# Numerical simulation and analysis of droplet formation within an amphiphilic particle

**DOI:** 10.1101/2023.10.11.561897

**Authors:** Xinpei Song, Ghulam Destgeer

## Abstract

An instrument-free particle-templated droplet formation can be achieved upon simple mixing of amphiphilic particles with aqueous and oil phases in a well plate by using a common lab pipette. Here, a two-dimensional, two-phase flow model was established using a finite element method to mimic the droplet formation within a concentric amphiphilic particle, which consisted of an outer hydrophobic layer and an inner hydrophilic layer. Immiscible water and oil phases selectively interacted with the hydrophilic and hydrophobic layers of the particle, respectively, to form an isolated aqueous compartment within a cavity. Three extreme models were also simulated, including completely hydrophilic, completely hydrophobic, and oppositely amphiphilic particle, which indicated that a right order of the particle layers was necessary to capture the droplet inside the cavity. Moreover, we performed a systematic study of particle-templated droplet formation by varying the individual layer thicknesses of particle, particle height, interfacial tension between water and oil, contact angle of interface with different surfaces, velocity of incoming oil media, and distance between neighboring particles. The volume fraction of water droplet trapped within the target cavity region was calculated to characterize the droplet formation. Our work will help to optimize the particle fabrication process, predict the experiment droplet formation, and explain the physical mechanism underlying compartmentalization phenomena.

## I. INTRODUCTION

Dispersion of a sample volume into numerous compartments containing target biomolecules or cells is a promising method to accumulate an amplifying signal to a detectable level. Uniform compartments are necessary to support similar reaction conditions for potential applications in biochemical sensing,^1^ nucleic acid detection,^2,3^ single cell analysis,^4,5^ drug discovery,^6^ and directed enzyme evolution.^7^ Microfluidic wells^8^ and droplet generators^9–11^ have been reported to form uniform compartments; however, these methods require expert users and specialized instruments.^12^ In contrast, particle-templated droplet or *dropicle* formation has been proposed as an instrument-free and user-friendly method to capture a uniform volume of aqueous solution inside or surrounding an engineered solid particle upon simple mixing with continuous oil and dispersed aqueous phases.^13–18^ Single-material hydrophilic particles, with a spherical or a crescent shape, hold a volume of an aqueous solution upon the shear-induced splitting of a larger aqueous volume.^19–21^ Multi-material amphiphilic particles of different shapes, composed of an inner hydrophilic layer and an outer hydrophobic layer, spontaneously capture a uniform droplet volume inside their cavities as the aqueous and oil phases prefer to interact with the inner hydrophilic and outer hydrophobic surfaces of the particle, respectively.^22–24^

The formation of uniform volume dropicles strongly depends upon the characteristics of the templating particles, e.g., particle dimensions, shape, and surface properties, in addition to the mixing conditions of the immiscible aqueous and oil phases. Experimentally varying these parameters to explore broader applications of the dropicles is costly and time-consuming. Numerical simulations of the dropicle formation provide a new prospect for revealing underlying mechanisms of droplet splitting based on various particle shapes and mixing conditions. Moreover, a numerical model simulating the dropicle formation would be indispensable in optimizing the microfluidic devices used for the particle fabrication with unique shapes and predicting the dropicle volume and uniformity, especially for irregular particle shapes. A mathematical model, based on minimizing the surface tension energy for a system of axisymmetric particles, has demonstrated that single material crescent or cylindrical-shaped particles can template uniform volume droplets.^25,26^ However, a computational fluid dynamic (CFD) simulation of a droplet captured by a solid particle, particularly with amphiphilic multi-material characteristics, is still needed to comprehend the underlying mechanism of droplet formation as it happens.

Here, we present a time-dependent CFD model based on a finite element method to simulate the formation of a droplet within a concentric amphiphilic particle. We developed a two-dimensional (2D) model comprising a cross-section of a bi-layered particle sitting on the bottom of a wellplate at the center of a multiphase fluidic domain. The particle captures an aqueous droplet within its hydrophilic cavity as an inflowing oil phase pushes the aqueous solution out of the fluidic domain and interacts with the outer hydrophobic layer of the particle. We performed a systematic study of dropicle formation within particles of variable heights, cavity diameters, and surface properties. We varied the thicknesses and contact angles for the hydrophilic and hydrophobic layers of the particle to investigate the uniform dropicle formation. Moreover, we analyzed the effect of medium density, interfacial tension, gravity, velocity of incoming oil phase, and distance between neighboring particles on the dropicle formation. The numerical model allowed us to simulate extreme cases of entirely hydrophilic or hydrophobic particles to highlight the need of an amphiphilic bi-layer particle for uniform dropicle formation.

## II. MODEL AND METHODS

Throughout this work, we have adopted a 2D model to mimic the droplet formation process within a 3D concentric amphiphilic microparticle, which has been experimentally studied before.^22,23^ **Figure 1**(a) depicts schematic diagram of the droplet formation process within the 3D particle, which was composed of an inner hydrophilic layer of polyethylene glycol diacrylate (PEGDA) and an outer hydrophobic layer of polypropylene glycol diacrylate (PPGDA). The particle was immersed in the water at the start. Then, the poly (dimethylsiloxane-co-diphenylsiloxane) (PSDS) oil was injected into the target region along the green arrow, pushing the water to flow towards the outlet on the right. As a result, an isolated droplet was formed inside the particle cavity as the excess water flowed away.

**Figure 1.**
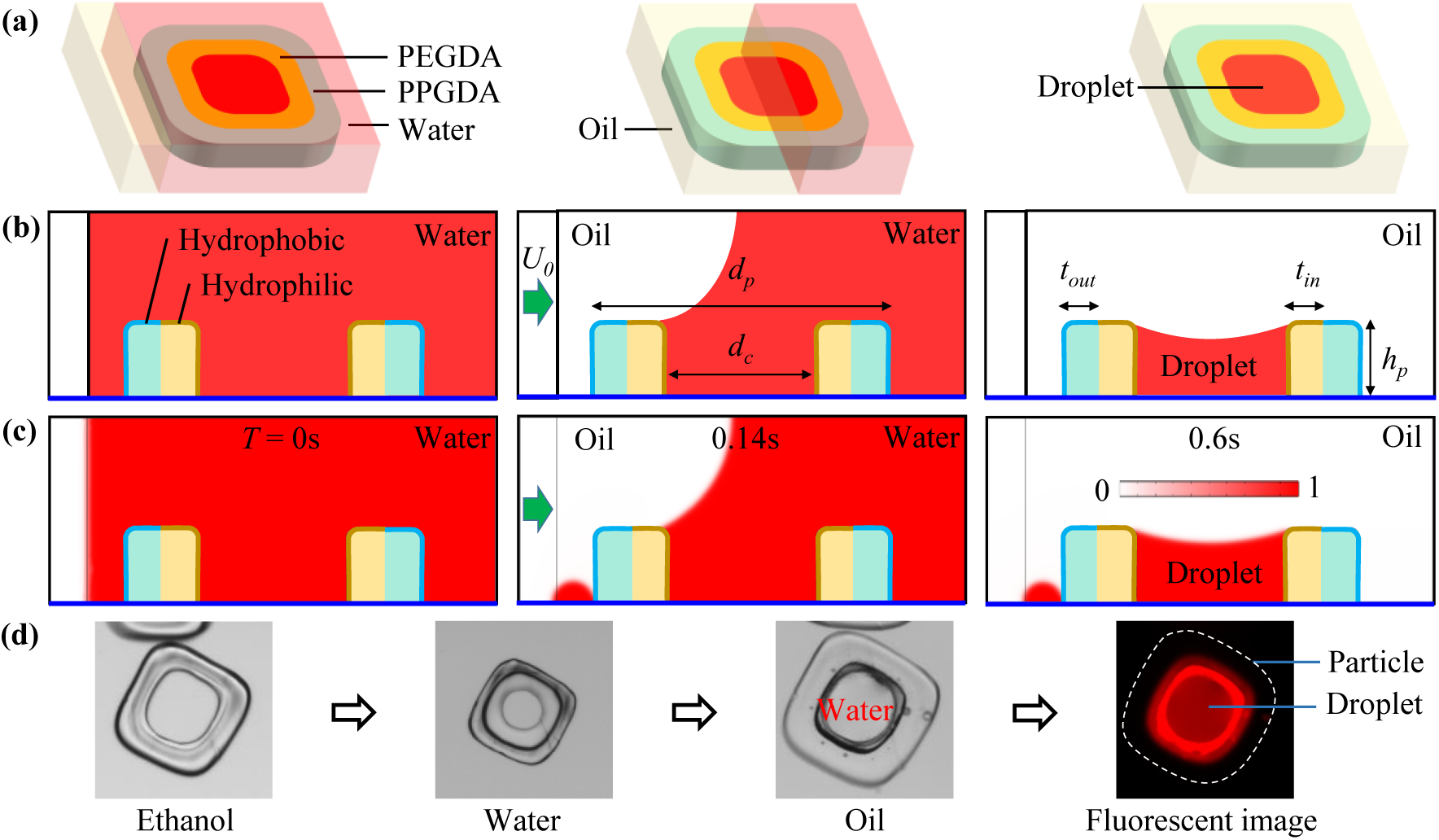
(a) Schematic diagram of a 3D concentric amphiphilic particle immersed in water. Injected oil into the domain pushes the excess water away to form a droplet. (b) Side view of the 3D schematic at three different stages. (c) A simulation of 2D model at *T*=0, 0.14s, and 0.6s. The red color legend represents the normalized volume fraction of water. (d) Experimental workflow of droplet formation within a particle.

The schematic in Figure 1(b) corresponds to the side view of the particle placed on a well plate at the three stages of the droplet formation. The highlighted brown and blue surfaces represent the inner hydrophilic and outer hydrophobic layers of the particle, respectively. The dark blue line at the bottom denotes the well plate surface.

To analyze the droplet formation within the particle, a 2D model was established using the two-phase flow module, containing the laminar flow and phase field, in COMSOL Multiphysics 6.1, a commercial software based on the finite element method. Figure 1(c) shows the volume fraction distribution of water and oil phases varying with the injection time of oil into the domain, i.e., *T* = 0s, 0.14s, and 0.6s. The width and height of the whole domain were 0.6mm and 0.25mm, respectively. The particle’s geometric parameters were as follows: height of particle *hp* = 0.1mm, diameter of particle *dp* = 0.4mm, diameter of particle cavity *dc* = 0.2mm, thickness of inner layer *tin* = 0.05mm, and thickness of outer layer *tout* = 0.05mm. Two liquids, water and oil, were adopted as the immiscible continuous phases. Water’s density and dynamic viscosity were 1000kg/m^3^ and 0.001Pa·s, and that of the oil were 1050kg/m^3^ and 0.06Pa·s, respectively. The interfacial tension between water and oil was *σ* = 0.03N/m. We continued injecting the oil for 0.55s with an average velocity of *U*0 = 1mm/s. The contact angles of the inner- and outer-layers of the particle with the water-oil interface were set as *θin* = 45° and *θout* = 150° in the simulations, corresponding to the hydrophilic and hydrophobic surface properties, respectively. The hydrophilic layer enabled the particle to capture a water droplet inside its cavity, which was isolated by the continuous oil phase interacting with the hydrophobic layer. The contact angle of the bottom well plate surface was *θwp*=90°. We employed an adaptive mesh, where a finer mesh region followed the water-oil interface as the simulation moved forward in time (see **Movie S1** and **Figure S1** in the supporting information). The thickness of the water-oil interface was set as 0.003mm to accurately trace the water-oil interface. The captured droplet was depicted in the form of volume fraction of water at *t*=0.6s.

Figure 1(d) shows the experimental workflow of the droplet formation within a particle.^23^ A fabricated particle was immersed in the ethanol, before the medium was exchanged to an aqueous solution mixed with a fluorescent dye, and finally oil was added to form a fluorescent droplet. The experimental result validates the numerical model.

## III. RESULTS AND DISCUSSIONS

### A. Extreme models

To explain the physical mechanism behind the dropicle formation within the amphiphilic particle, we established three extreme models with various particle surface properties by changing the contact angles of the inner- and outer-layer surfaces. The other parameters were identical to the basic model in Figure 1(c). In Figure 2(a), all the particle surfaces were hydrophobic, i.e., *θin* = *θout* = 150°. The water did not adhere to the particle, and an isolated dropicle was not formed covering the whole cavity of the particle. Two small satellite droplets were left behind on the well plate surface at *T* = 0.6s as the rapidly moving water-oil interface was broken at two locations due to the high shear rate at an injection velocity of *U*0 = 1mm/s. Figure 2(b) plots the simulation results when all the particle surfaces were hydrophilic, i.e., *θin* = *θout* =45°. The particle acted like a single-material hydrophilic particle, and both the inner and outer layers were surrounded by water.^27^ The dropicle formation inside the hydrophilic cavity was expected, whereas, in the absence of an outer hydrophobic layer, the oil flow was not able to remove the excess aqueous volume around the particle. In Figure 2(c), the particle possessed an opposite amphiphilic surface property compared to the basic model in Figure 1(c), i.e., inner layer was hydrophobic with *θin* = 150° and outer layer was hydrophilic with *θout* = 45°. An expected dropicle was not formed inside the cavity because the water preferred to adhere to the outer layers of the particle. Most of the water was pushed downstream of the particle due to high shear stress at the leading end of the particle, whereas, the trailing end of the particle was able to retain the aqueous media. By comparing the three models in Figure 2 with the basic counterpart in Figure 1(c), it was confirmed that the amphiphilic particle should have an inner hydrophilic layer and an outer hydrophobic layer for successful dropicle formation (see **Movie S2**).

**Figure 2.**
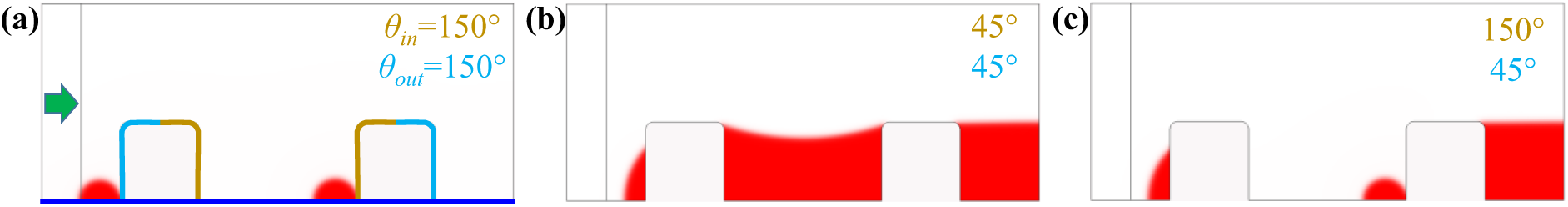
Extreme models of particle with variable surface properties. (a) All the particle surfaces were hydrophobic. (b) All the particle surfaces were hydrophilic. (c) The particle’s inner surface was hydrophobic and the outer surface was hydrophilic. The contact angle of the well plate surface was *θwp*=90°.

### B. Effects of surface properties on dropicle formation

#### 1. Contact angles of the particle and well plate surfaces

To figure out the effects of contact angles of the particle and well plate on the dropicle formation, we analyzed the model by varying *θin*, *θout*, and *θwp* in the range of 0-180° (see **Movie S3**). Figure 3(a) depicts the simulation results in the form of volume fraction of water with different *θin* of 0°, 90°, and 100°, while *θout* = 150° and *θwp* = 90°. The hydrophilicity of the inner-layer surface got weaker as we increased *θin* from 0 to 180°. The surface was perfectly hydrophilic when *θin* = 0, and a dropicle was formed inside the particle, with leading and trailing edges of the dropicle sitting at the boundaries between the inner and outer layers, as denoted by the dotted black lines. For a critical contact angle of *θin* = 90°, the inner particle surface was expected to change from hydrophilic to hydrophobic. Therefore, at *θin* = 90°, the dropicle volume within the cavity was significantly reduced from that with *θin* = 0°. With further increase in *θin* to 100°, the expected droplet was not formed because all the particle surfaces were hydrophobic at this point. To quantitatively analyze the effects of contact angle variation on the dropicle formation, we plotted the area of water in the whole domain (*SW*) normalized by the area of cavity (*S0*) at *T* = 0.6s by shifting *θin* from 0° to 180° with a step of 20° (Figure 3(b)). The normalized area (*SN* = *SW*/*S0*) of water represented the ratio between the water area in the target region and that of the cavity *S0* = 0.02mm^2^. If the expected dropicle was formed within the particle, the normalized area *SN* was close to 1. The normalized area was slightly lower than 1 when *θin*<90°, which meant that the expected droplet was formed. The *SN* value decreased gradually from ∼1 to ∼0.5 when increasing *θin* from 0 to 90°. The normalized area dips dramatically to <0.2 when *θin* > 90°, indicating that the expected droplet could not be formed. Here, *θin* = 90° was understood as the critical contact angle of the inner layer of the particle, below which an expected droplet was formed.

**Figure 3.**
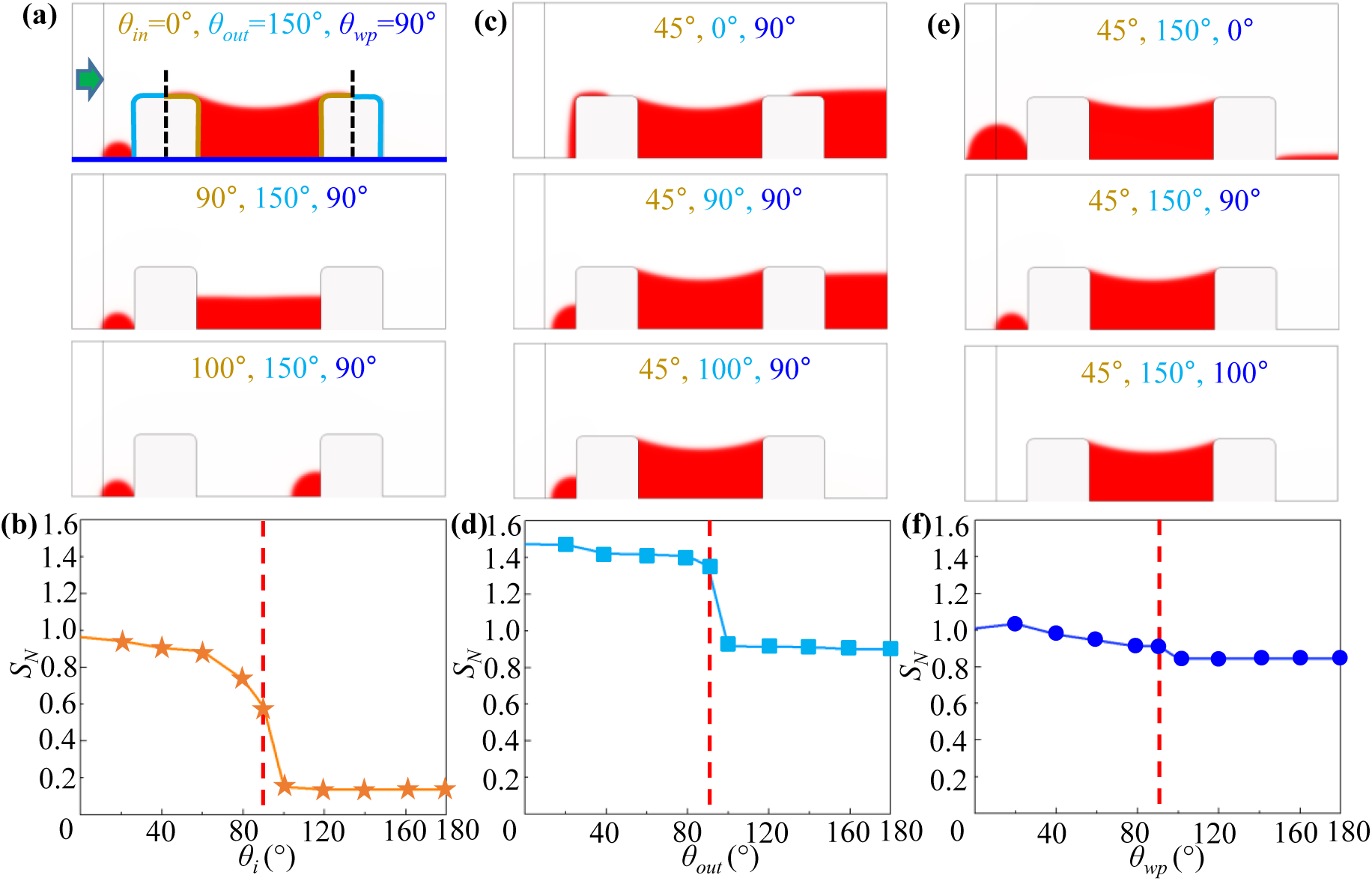
The effect of contact angles of the inner layer, outer layer, and well plate surface on the dropicle formation. (a) The volume fraction of water when *θin* = 0°, 90°, and 100° (a), *θout* = 0°, 90°, and100° (c), and *θwp* = 0°, 90°, 100° (e). Relationship between the normalized area of water *SN* and *θin* (b), *θout* (d), and *θwp* (e).

Similarly, in Figure 3(c), we plot the simulation results with different *θout* = 0°, 90°, and 100° to figure out the effects of the contact angle of the outer layer on the droplet formation, whereas *θin* = 45° and *θwp* = 90°. All surfaces of the particle were hydrophilic when *θout* was smaller than 90°. Therefore, the particle acted as a single-material hydrophilic particle, where the water adhered to both of the inner and outer layers of the particle. The expected droplet was formed because the droplet lacked the protection of an outer hydrophobic layer, and satellite droplets kept adhering to the outer layer of the particle. The amphiphilic property of the particle surfaces was restored when *θout* > 90°. Hence, an expected droplet was observed inside the cavity with *θout* = 100°. A critical contact angle of *θout* = 90° could be defined for the outer particle layer. We also plotted *SN* against increasing *θout* (Figure 3(d)). For *θout* < 90°, the hydrophilic surfaces of the particle captured a higher volume of water resulting in *SN* > 1. When *θout* > 90°, *SN* value approached 1, which meant that the expected droplet volume was captured by the particle.

Moreover, we varied the contact angle of the well plate surface *θwp* from 0 to 180°, where *θin* = 45° and *θout* = 150° (Figure 3(e)). It was observed that an expected dropicle was always formed within the particle irrespective of *θwp*, which was further confirmed by plotting *SN* against *θwp* (Figure 3(f)). The *SN* value floated around 1, indicating that the droplet formation was dominated by the amphiphilic property of the particle, and was not significantly affected by the well plate. However, the satellite droplets were easily removed from the domain when *θwp* > 90°. The inner hydrophilic inner layer was generally responsible for forming a droplet, and the hydrophobic outer layer had the capability of isolating the droplet.

#### 2. Two types of droplets depending on inner layer contact angle

To figure out the effect of the contact angle of the inner hydrophilic layer on the droplet shape, we simulated the volume fraction of water by adopting different *θin* (0° to 75°) in the basic model (Figure 1(c)), and keeping *θout* = 120° and *θwp* = 90° constant (see **Movie S4**). For super hydrophilic inner layer of the particle with *θin* = 0° and 15°, a droplet was always formed inside the cavity with an edge at the boundary between the inner and outer layers of the particle, as shown in Figure 4(a) and Figure 4(b), respectively. This type of droplet was labeled as Type Ⅰ. The super hydrophilic inner layer pulls the aqueous droplet as close as possible to wet all of the available hydrophilic surface, thereby, overcoming the interfacial tension pulling the droplet towards the center of the particle. Hence, the outer diameter of the droplet was defined by the thickness of the hydrophilic layer.

**Figure 4.**
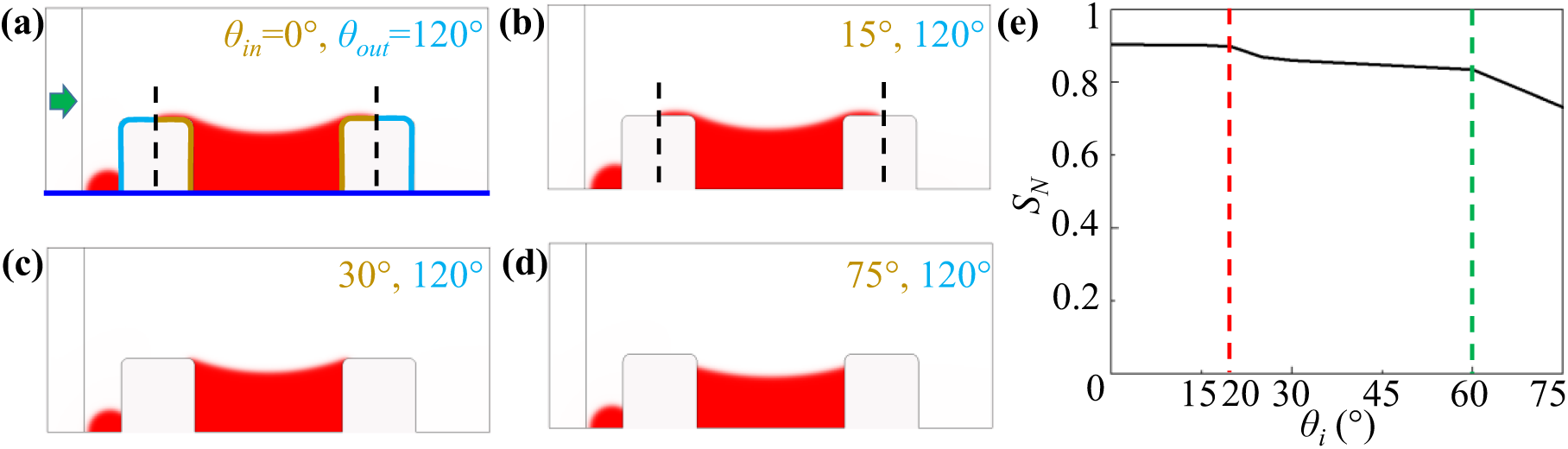
The effect of *θin* on the droplet type. Type Ⅰ droplet was formed with *θin* = 0 (a) and *θin*=15° (b). Type Ⅱ droplet was formed with *θin*=30° (c) and *θin*=45° (d). (e) Relation between the normalized area of the droplet *SN* and *θin*. *θout* = 120° and *θwp* = 90° were constant.

The edge of the droplet dropped inside the cavity when the inner layer contact angle increased to *θin* = 30° and 75°, as depicted in Figures 4(c) and 4(d), respectively. We marked this kind of droplet as Type Ⅱ. The interfacial tension was able to pull the droplet away from the top surface of the particle towards the center since the wettability of the hydrophilic surface dropped with the increase in contact angle. We plotted *SN* as *θin* was increased in small steps from 0° to 75° (Figure 4(e)). For *θin* < 20°, *SN* was slightly higher compared to other simulations when *θin* > 20° because of the droplet stretching on top of the particle. The droplet area decreased significantly when *θin* > 60°. However, it was noted that the droplet diameter would not be affected by the inner layer thickness *tin*, when *θin* > 20° since the water-oil interface will be interacting within the inner surface of the cavity.

#### 3. Effect of particle symmetry on dropicle formation

We used Type Ⅰ droplets to study the effects of the particle symmetry on the droplet formation by adopting *θin* = 18° and *θout* = 162° (see **Movie S5** and Figure 5(a)). The symmetrical and asymmetrical particles were designed by adjusting the thicknesses of the inner and outer layers, *tin* and *tout*, while *tin* + *tout* = 0.1mm was kept constant. Figure 5(a) depicts the water volume fraction in the whole domain after the droplet was formed using symmetrical particles with *tin* = 0.08mm, 0.07mm, 0.04mm, and 0.02mm, respectively. The expected droplet was not formed when *tin* = 0.08mm because the outer hydrophobic layer thickness (*tout* = 0.02mm) was too small to break the water-oil interface at the trailing edge of the particle under the given inlet velocity. As *tin* was reduced, a symmetric Type Ⅰ droplet was observed with the edge at the boundary between the inner and outer layers of the symmetric particle. The *SN* value was much higher than 1 with *tin* = 0.08mm, indicating that a large volume of water was captured by the particle (Figure 5(b)). The *SN* value approached 1 when *tin* < 0.08mm.

**Figure 5.**
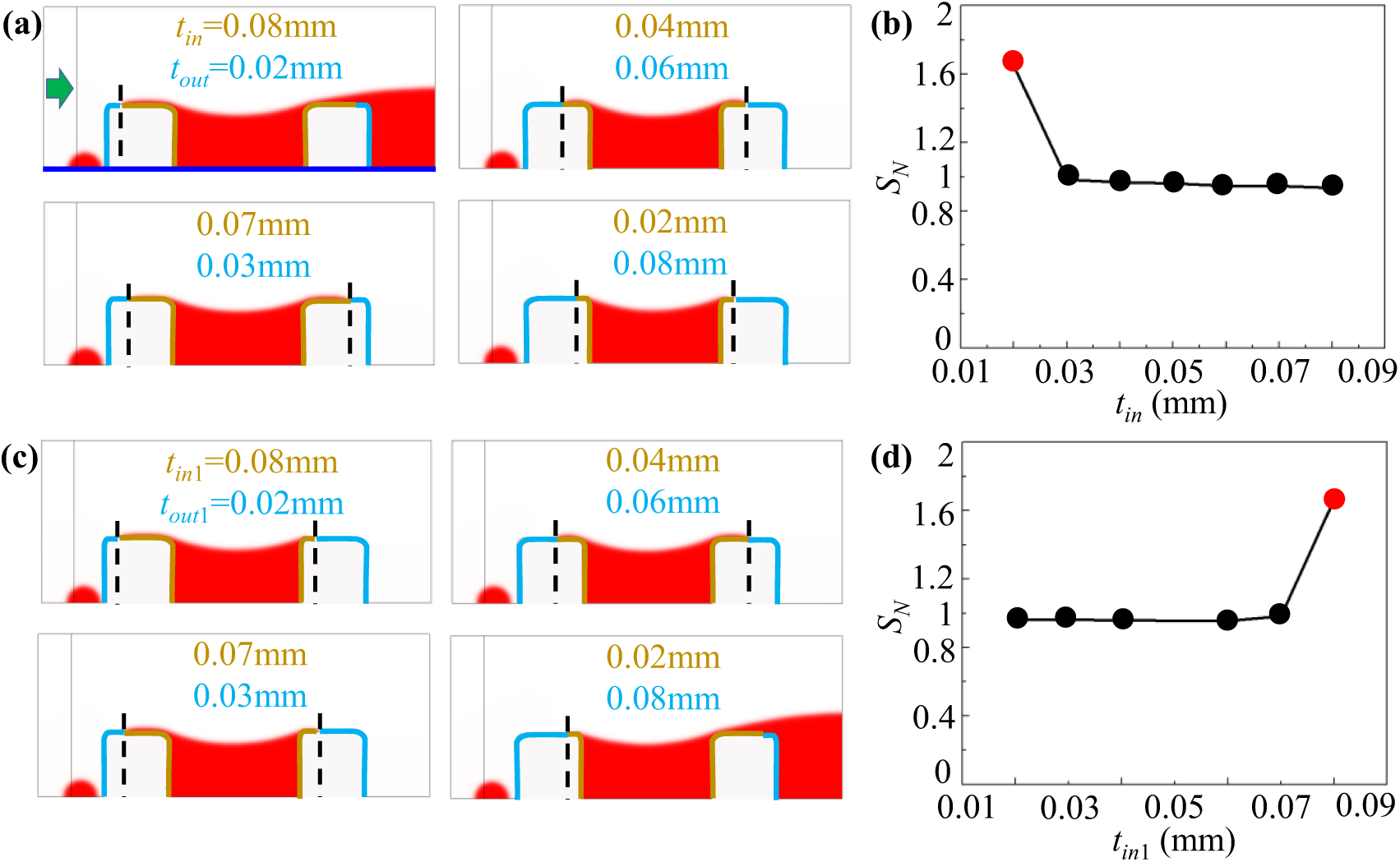
The effect of symmetric and asymmetric particles on the dropicle formation. (a) Volume fraction of water for the symmetric particle with *tin*=0.08mm, 0.07mm, 0.04mm, and 0.02mm. (b) Normalized area of water (*SN*) for the symmetric particle with various *tin*. (c) Volume fraction of water for the asymmetric particle with *tin*1=0.08mm, 0.07mm, 0.04mm, and 0.02mm. (d) *SN* for the asymmetric particle with various *tin*1.

For an asymmetric particle, the thicknesses of the inner and outer layers of the left section of the particle were defined as *tin*1 and *tout*1, and those of the right section were *tin*2 and *tout*2, respectively. Here, *tin*1 = *tout*2, *tin*2 = *tout*1, *tin*1 + *tout*1 = 0.1mm, and *tin*2 + *tout*2 = 0.1mm. Figure 5(c) shows the simulation results in the form of water volume fraction when *tin*1 = 0.08mm, 0.07mm, 0.04mm, and 0.02mm, respectively. The asymmetric droplets were formed within the asymmetric particles for the former three cases. The *SN* value was >1 when *tin*1 = *tout*2 = 0.02mm because the thin outer hydrophobic layer at the trailing edge of the particle could not break the water-oil interface (Figure 5(d)). It was observed that the larger thickness of the outer hydrophobic layer, particularly at the trailing end, was critical to breaking the water-oil interface and capturing a droplet within the cavity.

#### 4. Effect of oil velocity on dropicle formation

To analyze the effects of the inlet velocity of oil *U0* on the droplet formation, we used the basic model (Figure 1(c)) to simulate the volume fraction of water by shifting *U0* from 0.1mm/s to 20mm/s. The selected results show a Type Ⅱ droplet inside the cavity when *U0*=0.1mm/s, 2mm/s, and 20mm/s, respectively (see **Movie S6** and Figure 6(a)). When *U0*=0.1mm/s, an expected dropicle was observed without extra satellite droplets because the slow inlet velocity yielded enough time to push away the excess water outside of the particle cavity before the water-oil interface could break. The satellite droplets emerged when increasing *U0* to 20mm/s, but it had little effect on the dropicle formation within the cavity. The droplet formation time was shortened as the inlet velocity *U0* was increased from 0.1mm/s to 20mm/s (Figure 6(b)). The dropicle formation time decreased sharply when *U0* changed from 0 to 2mm/s, and decreased slowly for *U0* > 2mm/s. The formation time could be dramatically shortened by properly increasing the injecting velocity of oil, without significantly effecting the dropicle shape.

**Figure 6.**
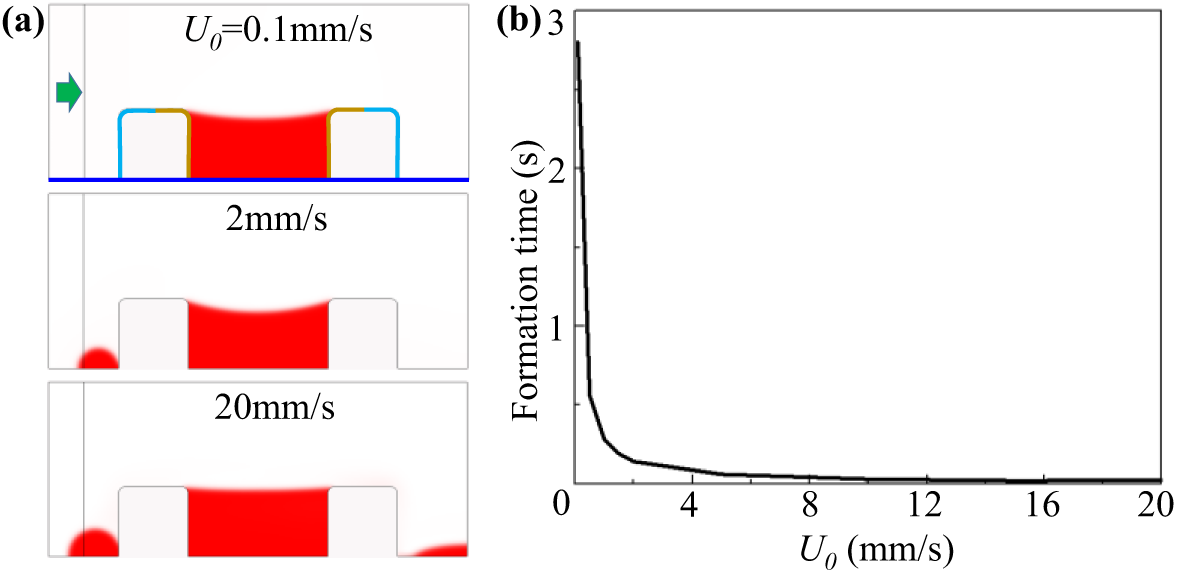
The effect of inlet velocity on the dropicle formation. (a) The volume fraction of water when *U0*=0.1mm/s, 2mm/s, and 20mm/s. (b) Relation between the dropicle formation time and *U0*.

#### 5. Effect of interfacial tension on dropicle formation

The effect of interfacial tension *σ* on the droplet formation was analyzed using the basic model (Figure 1(c)). The water volume fraction was simulated when *σ* = 0N/m, 0.001N/m, and 0.1N/m, respectively (see **Movie S7** and Figure 7(a)). A dropicle usually forms when the shear-induced drag force takes away the excess aqueous volume with the oil flow, whereas the interfacial tension retains a droplet within the hydrophilic cavity of the particle. Therefore, when the interfacial tension was ignored, i.e., *σ* = 0N/m, while the shear-induced drag force was still active, no dropicle was captured. However, a dropicle was formed with the introduction of a small interfacial tension of *σ* = 0.001N/m. Further increase in *σ* to 0.03N/m and 0.1N/m resulted in dropicle formation inside the cavity. We plotted *SN* by shifting *σ* from 0 to 0.1N/m (Figure 7(b)). The *SN* value > 1 when *σ* = 0N/m, and the expected droplet was not formed. However, the *SN* value approaches 1 after introducing *σ* to the model.

**Figure 7.**
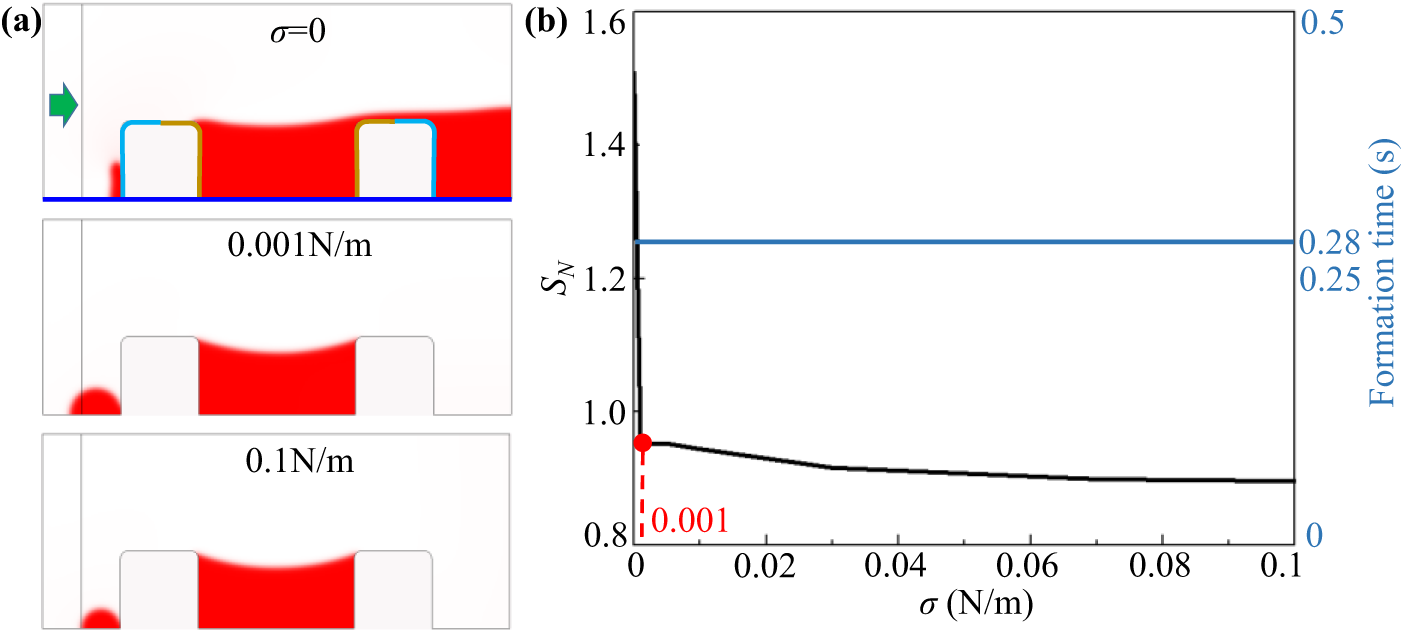
The effect of interfacial tension on the dropicle formation. (a) Volume fraction of water under diverse interfacial tension *σ* = 0, 0.001N/m, and 0.1N/m. (b) The *SN* (black line) and formation time (blue line) with variable *σ*.

#### 6. Other parameters and properties

The effects of other geometric parameters of the particle were also analyzed using the basic model (Figure 1(c)), including the height of particle *hp* (**Figure S2**) and diameter of particle cavity *dc* (**Figure S3**). The droplet could not be captured inside the cavity when *hp* was too short. Dropicles were captured within particles of varying *dc*. The density of oil (**Figure S4**) and gravity (**Figure S5**) did not dominate the droplet formation.

### C. Interaction between neighboring particles

To understand the interaction between neighboring particles, we simulated the volume fraction of water to observe the droplet formation, and we also compared the normalized area (*SN*) of each droplet within the target region (Figure 8). Figure 8 shows the results for Type Ⅰ droplets, and that for Type Ⅱ droplets are depicted in **Figure S6**. For the Type Ⅰ droplet, the model comprised of two identical amphiphilic particles with *θin* = 18°, *θout* = 162°, and *θwp* = 90°. The distance between the particles was *D*. Here, we adopted *D* = 0.01mm, 0.1mm, and 0.2mm to plot the volume fraction of water, respectively (Figure 8(a)). We observed four isolated droplets in the whole domain, two of which were dropicles within the designated particle cavities with *SN* → 1, the other two were satellite droplets with *SN* << 1 for different values of *D* (Figure 8(b)). The two uniform droplets within two identical amphiphilic particles were not affected by the distance between the neighboring particles. Their shapes were similar, and the normalized areas were close to each other.

**Figure 8.**
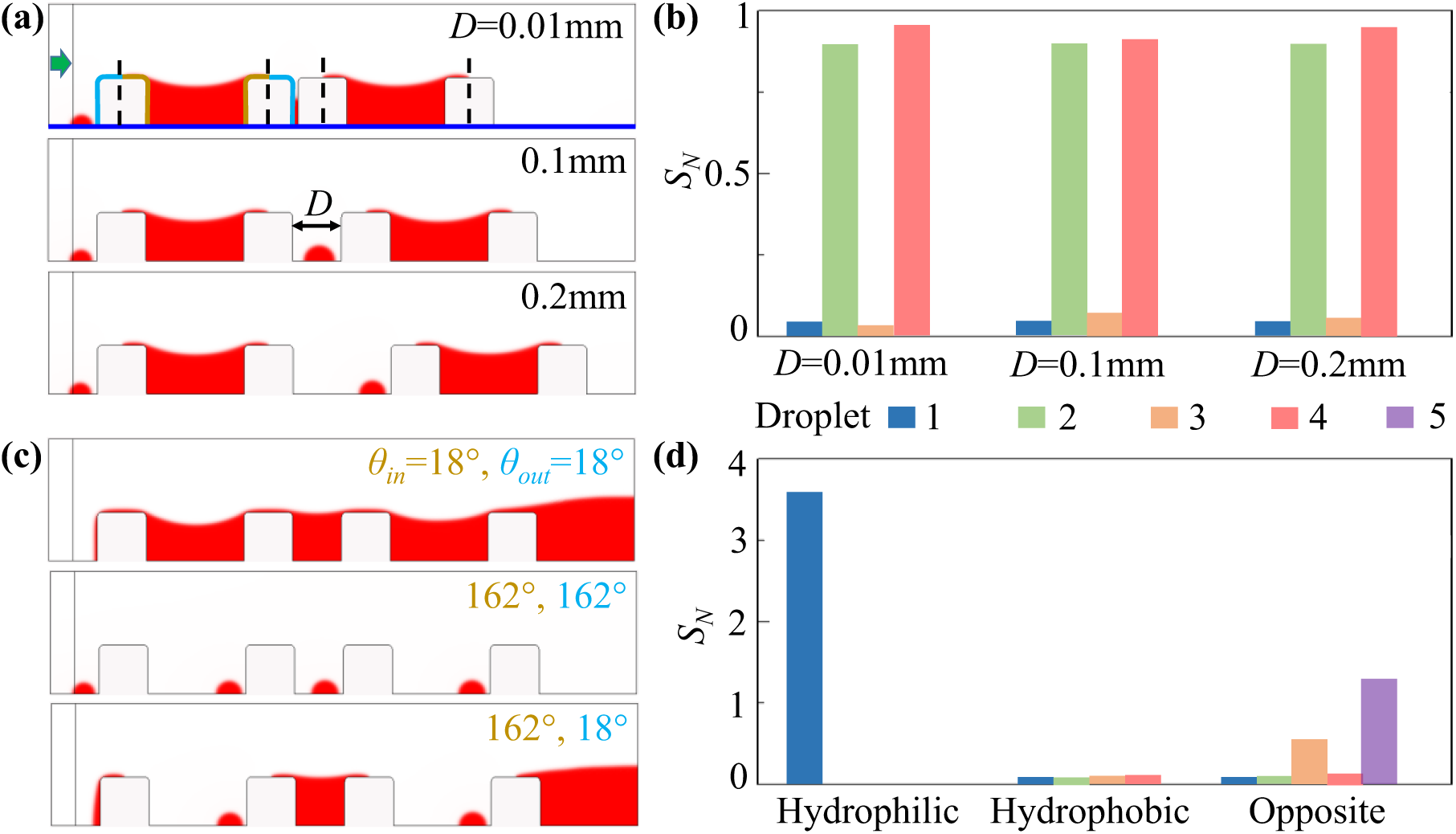
Interaction between neighboring particles. (a) Volume fraction of water with the two amphiphilic particles when interparticle distance *D* = 0.01mm, 0.1mm, and 0.2mm, respectively. (b) Normalized area of each droplet (1-g from left to right in (a)) with different *D*. (c) Volume fraction of water with *D* = 0.1mm in the extreme models that all surfaces of the particles are hydrophilic, all surfaces of the particles are hydrophobic, and the particle surface has the oppositely amphiphilic property to the basic model (Figure 2). (d) Normalized area of each droplet (1-5 from left to right) in the extreme models in (c), respectively.

We used the three extreme models containing two identical particles of hydrophilic, hydrophobic, and oppositely amphiphilic properties, as shown in Figures 8(c) and 8(d), respectively (also depicted earlier in Figure 2). The inter-particle distance *D* = 0.1mm was fixed. For the hydrophilic particles, *θin* = *θout* =18°. The two particles were surrounded by a single large water droplet. For the hydrophobic particles, *θin* = *θout* = 162°, four small satellite droplets were dispersed within the fluid domain. For the oppositely amphiphilic particles, *θin* = 162° and *θout* = 18°, we observed five droplets with non-uniform normalized area (*SN*). The water droplets were trapped between the neighboring particles instead of within the cavities. Figure 8(d) plots the *SN* for the three cases, respectively. Only a single bar for hydrophilic particles was noted with *SN* = 3.63, depicting one large droplet around the particles. The second set of bars shows that four droplets formed had *SN* << 1, indicating satellite droplet formation. For the particles possessing the opposite amphiphilic property, five non-uniform droplets were obtained. Overall, we did not obtain the expected droplets within these three classes of the particle.

## IV. CONCLUSIONS

A 2D model mimicking droplet formation within a concentric 3D amphiphilic particle has been established using the COMSOL Multiphysics software. A systematical work was conducted by studying three extreme models of particles, the effects of their surface properties on dropicle formation, and the interaction between neighboring particles. The dropicle formation was attributed to the amphiphilic properties of the particle surfaces, which was confirmed by analyzing the three extreme models. The effects of the contact angles of the inner and outer layers of the particle and the bottom well plate surface on the droplet formation were studied in detail, showing that the wetting behavior of the particle layers was critically important for the dropicle formation. Two different types of droplets were identified by tuning the contact angle of the inner layer of the particle. The thicknesses of the inner and outer layers were varied in a symmetrical and non-symmetrical manner to conclude that the outer hydrophobic layer thickness was significantly important in breaking the water-oil interface and forming isolated dropicles. We have further analyzed the effects of oil velocity, interfacial tension, height of particle, diameter of cavity, oil density, and gravitational force on dropicle formation. Finally, the interaction between neighboring particles was studied by establishing a model containing two identical particles. The uniform droplets were formed within two identical neighboring particles with amphiphilic properties. This numerical work covers a majority of the possibilities of droplet formation within a concentric amphiphilic particle, guiding the design and fabrication of new microfluidic devices at the heart of our experimental setup, significantly reducing the time and costs associated with a trial-n-error-based workflow. However, the presented 2D numerical models are not meant to completely replace the 3D models validated by experimental work.

## ACKNOWLEDGEMENT

X.S. gratefully acknowledges the Sino-German (CSC-DAAD) Postdoc Scholarship.

## Supporting Information

In this work, an adaptive mesh was employed in the model. Thirty meshes were generated in the simulation process, and each mesh moved following the water-oil interface. **Figure S1** shows several selected meshes. The first mesh highlighted the water-oil interface with a very refined mesh at the inlet of the domain. The sixteenth mesh corresponded to the droplet formation inside the cavity. Then, the extra water flowed away from the fluidic domain. Finally, an expected droplet was formed within the particle in the last frame.

**Figure S1.**
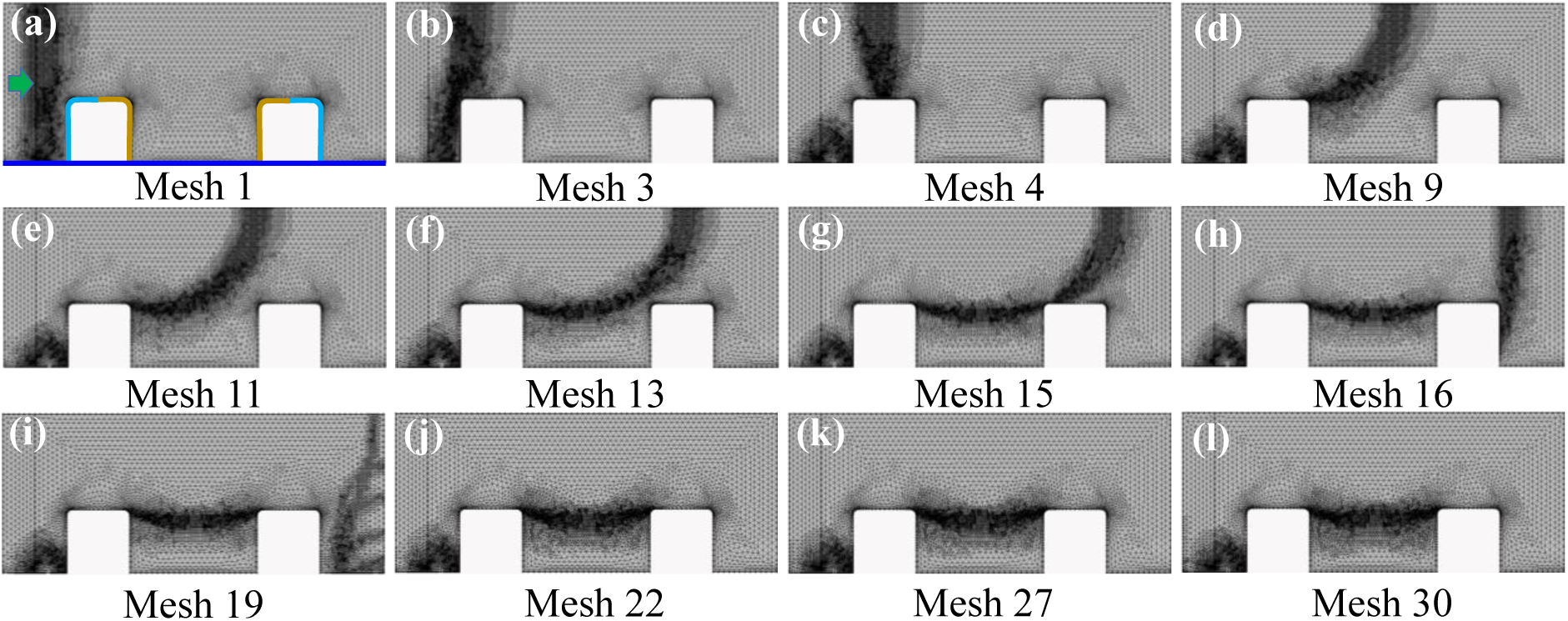
The adaptive meshes in the simulation process.

The effects of the geometric parameters of the particle, oil density, and gravity on the droplet formation were analyzed using the basic numerical model. The effects of geometric parameters of particles on droplet formation were studied by shifting the particle height *hp* and diameter of particle cavity *dc* in a reasonable range. **Figure S2** plots four selected simulation results when shifting *hp* from 0.01mm to 0.2mm. No droplet was obtained when *hp* = 0.01mm, as shown in Figure S2(a). The water was not captured by the particle because the height of the particle was too short to hold a droplet inside the cavity. Figures S2(b)-S2(d) plot the simulation results with *hp* = 0.03mm, 0.05mm, and 0.2mm, respectively. A convex-shaped droplet was formed inside the tallest cavity.

**Figure S2.**
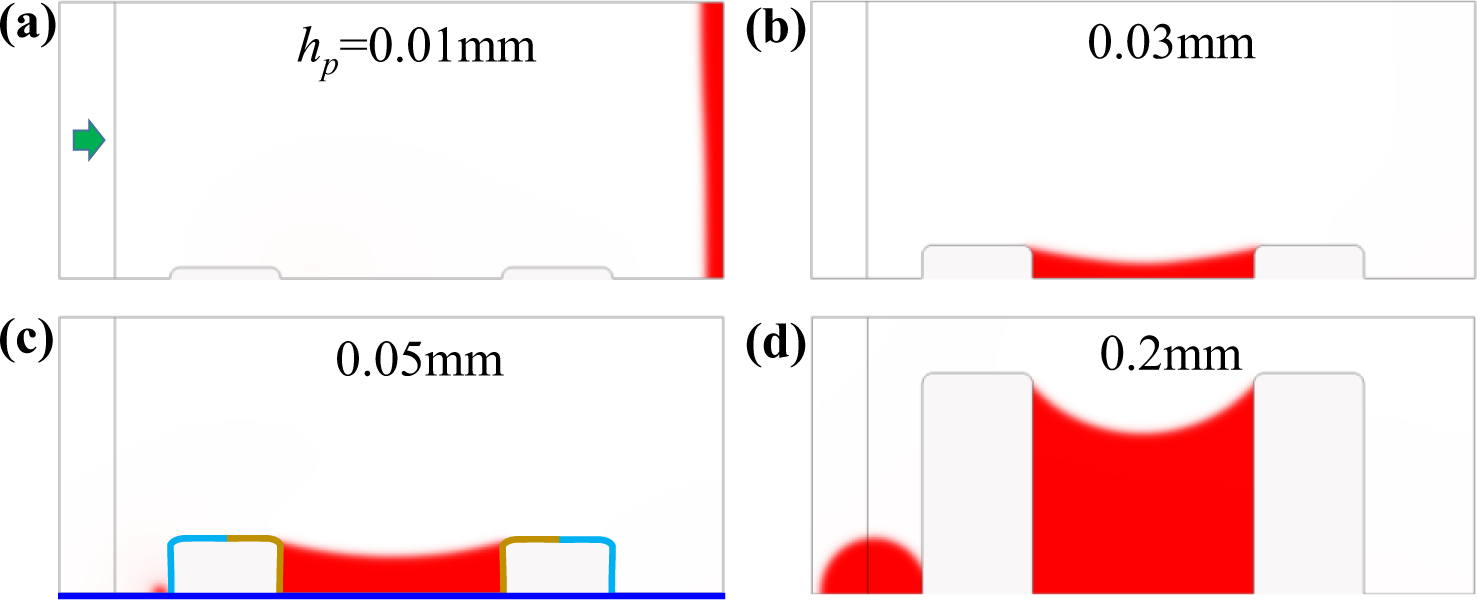
Effects of the particle height. (a) *hp*=0.01mm, (b) *hp*=0.03mm, (c) *hp*=0.05mm, and (d) *hp*=0.2mm.

**Figure S3.**
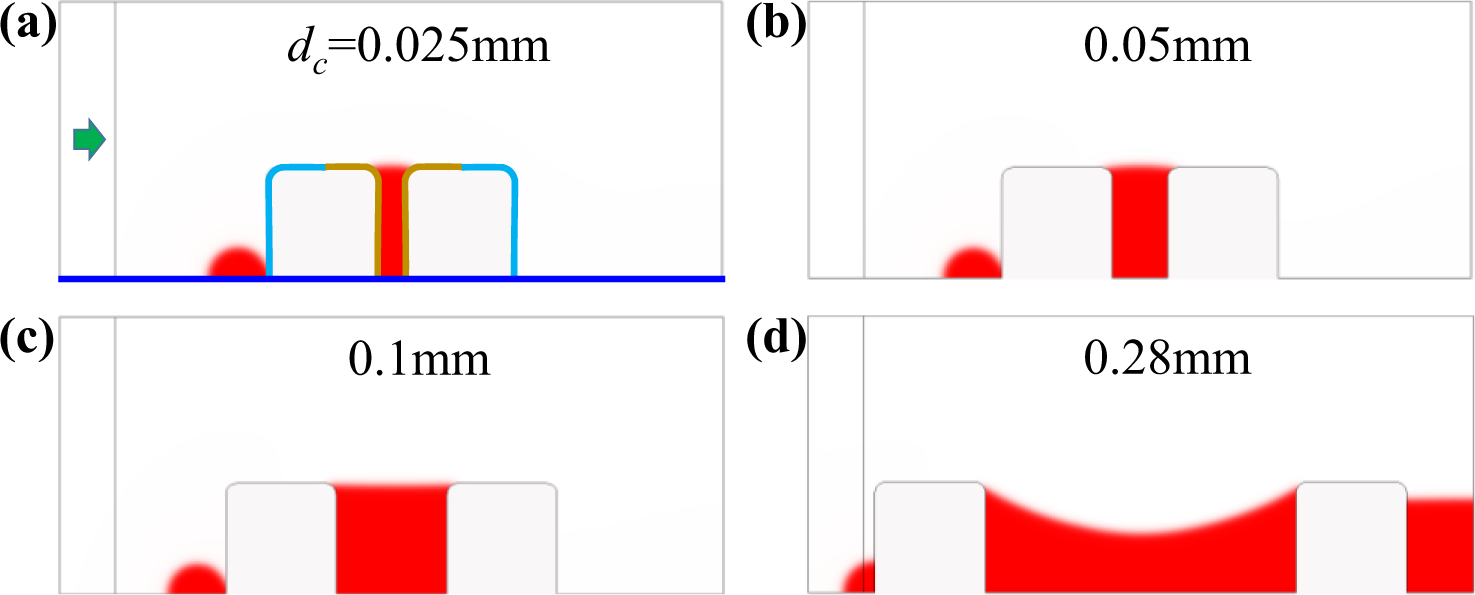
Effects of the diameter of particle cavity. (a) *dc*=0.025mm, (b) *dc*=0.05mm, (c) *dc*=0.1mm, and (d) *dc*=0.28mm.

Figure S3 presents the simulation results with *dc* = 0.025mm, 0.05mm, 0.1mm, and 0.28mm, respectively. The shape of the droplet surface was affected by *dc*. The droplet had a convex surface when *dc* > 0.1mm because the wide cavity enabled the shear-induced removal of aqueous volume from the cavity.

The simulations were conducted to figure out the effects of oil density *ρ* on the droplet formation by shifting *ρ* from 900kg/m_3_ to 1200kg/m^3^. The selected results are depicted in **Figures. S4**(a) and S4(b) for *ρ*=900kg/m^3^ and *ρ*=1200kg/m^3^, respectively. The formed droplets are similar to that when *ρ*=1050kg/m^3^, indicating that the oil density hardly affects the droplet formation.

**Figure S4.**
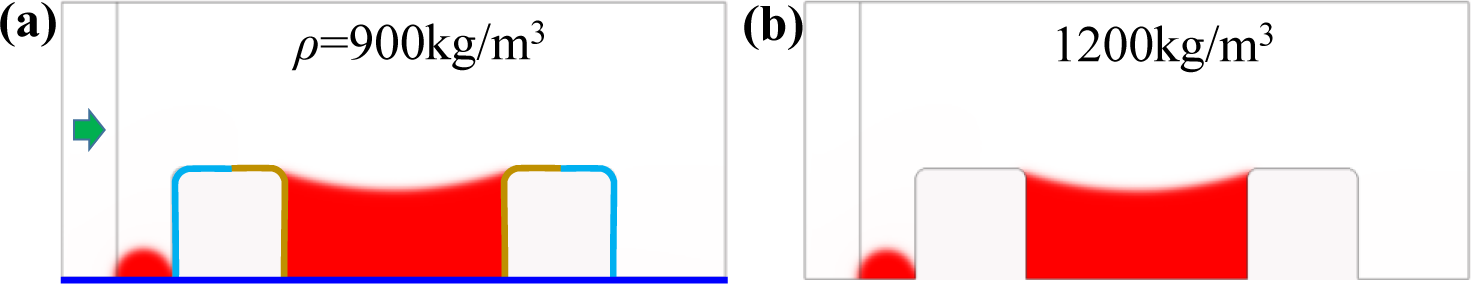
Effects of oil density. (a) *ρ*=900kg/m^3^ and (b) *ρ*=1200kg/m^3^.

The effect of gravity on the droplet formation was analyzed. The results are shown in **Figure S5**. The results without consideration of gravity were similar to those when gravity was considered. Thus, gravity was not a key factor in droplet formation at the current scale of particle height.

**Figure S5.**
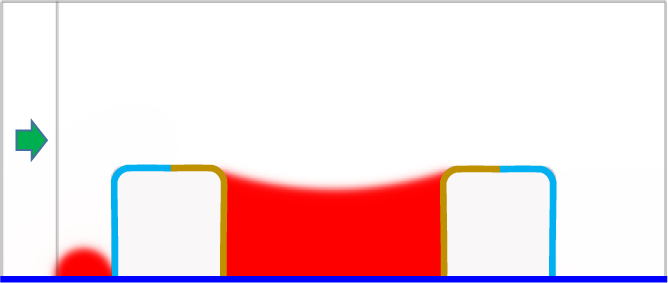
Effect of gravity on dropicle formation.

Similar to the analysis of Type Ⅰ droplet formation, the interaction between neighboring particles for Type Ⅱ droplet formation was analyzed qualitatively and quantitatively. In **Figure S6**(a), we plot the water volume fraction for the amphiphilic particles with three different distances, *D* = 0.01mm, 0.1mm, and 0.2mm. Here, *θin* = 45°, *θout* = 150°, and *θwp* = 90°. Four different droplets were seen in the whole region. The corresponding normalized areas of the four droplets are shown in Figure S6(b). *SN1* = 0.053, *SN2* = 0.853, *SN3* = 0.032, and *SN4* = 0.872, when *D* = 0.01mm. Similarly, *SN1* = 0.054, *SN2* = 0.851, *SN3* = 0.087, and *SN4* = 0.847, when *D*=0.1mm. *SN1* = 0.053, *SN2* = 0.851, *SN3* = 0.071, and *SN4* =0.870, when *D*=0.2mm. *SN2* and *SN4* were close to 1, and *SN1* and *SN3* were much smaller than 1. It indicates that the second and fourth droplets were as expected. Both the qualitative and quantitative results were similar to that for Type Ⅰ droplet formation. Two uniform droplets were formed within two identical amphiphilic particles, and the droplet formation in each particle was rarely affected by its neighboring particles.

Figure 6S(c) shows three models containing two identical particles of hydrophilic, hydrophobic, and oppositely amphiphilic properties individually. *D* = 0.1mm was fixed. When *θin* = *θout* = 45°, all the particle surfaces were hydrophilic, and we observed five droplets in the fluidic domain. Four small droplets were observed when *θin* = *θout* = 150°, meaning that all the particle surfaces were hydrophobic. The particle possessed the oppositely amphiphilic surface property when *θin*=150° and *θout*=45°, forming five different droplets. Hence, we could not obtain the expected droplet inside the particle cavities using these three models. Figure 6S(d) shows the normalized areas of each droplet for the three models, respectively. Five droplets were formed for the particles with all surfaces hydrophilic, *SN1* = 0.063 and *SN3* = 0.500 are much smaller than 1, indicating that no droplets were formed. *SN2* = 0.846 and *SN4* = 0.849 were close to 1, but the droplets are vulnerable to the water surrounding the outer layers of the two particles. *SN5* = 1.137 was close to 1 because the water surrounded the outer layer of the particle on the right side. Hence, the expected droplets were not observed. Four droplets were formed for the particles with all surfaces hydrophobic, and their normalized areas were *SN1* = 0.054, *SN2* = 0.070, *SN3* = 0.077, and *SN4* = 0.077. No expected droplets were formed. For the particles possessing the oppositely amphiphilic surface property, five droplets were obtained. *SN1* = 0.063, *SN2* = 0.070, *SN3* = 0.503, and *SN4* = 0.087 were much smaller than 1. The normalized area of the fifth droplet *SN5* =1.113 was close to 1, but it was outside of the particle cavity, meaning that we cannot get separated droplets within the particles.

**Figure S6.**
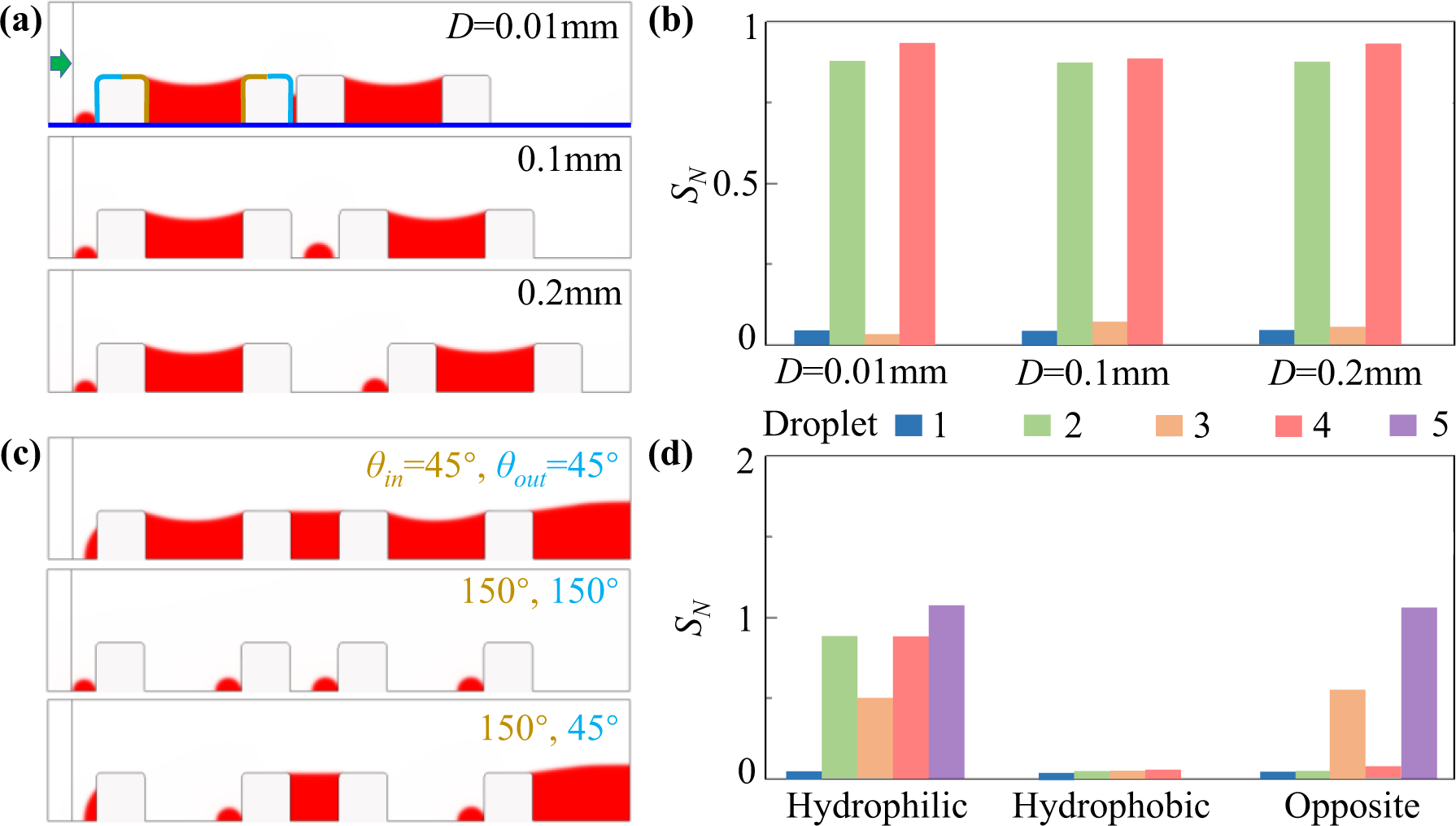
Interaction between neighboring particles. (a) Volume fraction of water for the amphiphilic particles with *D*=0.01mm, 0.1mm, and 0.2mm. (b) Normalized area of every droplet formed in (a), respectively. (c) Volume fraction of water with *D*=0.2mm for the extreme models that all surfaces of the particles are hydrophilic, all surfaces of the particles are hydrophobic, and the particle surface has the oppositely amphiphilic property to the basic model. (d) Normalized area of every droplet formed in (c), respectively.

## Movies captions

**Movie S1:** The adaptive mesh moves with the water-oil interface as the expected droplet is captured within the cavity.

**Movie S2:** Dropicle formation within extreme models of particles with variable surface properties corresponding to Figure 2.

**Movie S3:** The effect of contact angles of the inner layer, outer layer, and well plate surface on the dropicle formation corresponding to Figure 3.

**Movie S4:** The effect of *θin* on the droplet type as depicted in Figure 4.

**Movie S5:** The effect of symmetric and asymmetric particles on the dropicle formation as depicted in Figure 5.

**Movie S6:** The effect of inlet velocity on the dropicle formation as depicted in Figure 6.

**Movie S7:** The effect of interfacial tension on the dropicle formation as depicted in Figure 7.

**Movie S8:** The effect of neighboring particles on the dropicle formation as depicted in Figure 8.

## REFERENCES

1. I. Rea, E. Orabona, A. Lamberti, I. Rendina, and L. D. Stefano, “A microfluidics assisted porous silicon array for optical label-free biochemical sensing,” Biomicrofluidics 5, 034120 (2011).

2. X. Li, X. Zhao, W. Yang, F. Xu, B. Chen, J. Peng, J. Huang, and S. Mi, “Stretch-driven microfluidic chip for nucleic acid detection,” Biotechnol. Bioeng. 118, 3559–3568 (2021).

3. H. Lu, J. Zhu, T. Zhang, X. Zhang, X. Chen, W. Zhao, Y. Yao, W. Zhao, and G. Sui, “A rapid multiplex nucleic acid detection system of airborne fungi by an integrated DNA release device and microfluidic chip,” Talanta 246, 123467 (2022).

4. K. Matula, F. Rivello, and W. T. S. Huck, “Single-cell analysis using droplet microfluidics,” Adv. Biosyst. 4, e1900188 (2020).

5. J. D. Rutte, R. Dimatteo, M. M. Archang, M. V. Zee, D. Koo, S. Lee, A. C. Sharrow, P. J. Krohl, M. P. Mellody, S. Zhu, J. Eichenbaum, M. K, S. Udani, K. Ha, A. L. Bertozzi, J. B. Spangler, R. Damoiseaux, and D. D. Carlo, “Suspendable hydrogel nanovials for massively parallel single-cell functional analysis and sorting,” ACS Nano 16, 7242–7257 (2022).

6. F. He, M. J. Zhang, W. Wang, Q. W. Cai, Y. Y. Su, Z. Liu, Y. Faraj, X. J. Ju, R. Xie, and L. Y. Chu, “Designable polymeric microparticles from droplet microfluidics for controlled drug release,” Adv. Mater. Technol-US 4, 1800687 (2019).

7 A. Stucki, J. Vallapurackal, T. R. Ward, and P. S. Dittrich, “Droplet microfluidics and directed evolution of enzymes: an intertwined journey,” Angew. Chem. Int. Ed. 60, 24368–24387 (2021).

8. R. J. Shilton, L. Y. Yeo, and J. R. Friend, “Quantification of surface acoustic wave induced chaotic mixing-flows in microfluidic wells,” Sensor. Actuat. B-Chem. 160, 1565–1572 (2011).

9. R. Lathia, K. N. Nampoothiri, N. Sagar, S. Bansal, C. D. Modak, and P. Sen, “Advances in microscale droplet generation and manipulation,” Langmuir 39, 2461–2482 (2023).

10. F. M. Galogahi, A. Ansari, A. J. T. Teo, H. Cha, H. An, and N. T. Nguyen, “Fabrication and characterization of core-shell microparticles containing an aqueous core,” Biomed. Microdevices 24, 1–10 (2022).

11. T. Franke, A. R. Abate, D. A. Weitz, and A. Wixforth, “Surface acoustic wave (SAW) directed droplet flow in microfluidics for PDMS devices,” Lab Chip 9, 2625–2627 (2009).

12. V. Yelleswarapu, J. R. Buserb, M. Habera, J. Barona, E. Inapuria, and D. Issadore, “Mobile platform for rapid sub–picogram-per-milliliter, multiplexed, digital droplet detection of proteins,” Proc. Natl. Acad, Sci. U.S.A. 116, 4489–4495 (2019).

13. R. Novak, Y. Zeng, J. Shuga, G. Venugopalan, D. A. Fletcher, M. T. Smith, and R. A. Mathies, “Single-cell multiplex gene detection and sequencing with microfluidically generated agarose emulsions,” Angew. Chem. Int. Ed. 50, 390–395 (2011).

14. M. N. Hatori, S. C. Kim, and A. R. Abate, “Particle-templated emulsification for microfluidics-free digital biology,” Anal. Chem. 90, 9813–9820 (2018).

15. M. A. Sahin, M. Shehzad, and G. Destgeer, “Stopping microfluidic flow,” arXiv:2308.02386 [physics.flu-dyn].

16. V. Shah, X. Yang, A. Arnheim, S. Udani, D. Tseng, Y. Luo, M. Ouyang, G. Destgeer, O. Garner, H. Koydemir, A. Ozcan, and D. D. Carlo, “Amphiphilic particle-stabilized nanoliter droplet reactors with a multi-modal portable reader for distributive biomarker quantification,” doi: 10.1101/2023.04.24.538181.

17. M. U. Akhtar, M. A. Sahin, H. Werner, and G. Destgeer, “Fabrication of size-coded amphiphilic particles with a configurable 3D-printed microfluidic device for the formation of particle-templated droplets,” doi: 10.1101/2023.09.20.558669.

18. M. A. Sahin, H. Werner, S. Udani, D. D. Carlo, and G. Destgeer, “Flow lithography for structured microparticles: fundamentals, methods and applications,” Lab Chip 22, 4007 (2022).

19. Q. Liu, M. Zhao, S. Mytnyk, B. Klemm, K. Zhang, Y. Wang, D. Yan, E. Mendes, and J. H. V. Esch, “Self-orienting hydrogel micro-buckets as novel cell carriers,” Angew. Chem. 131, 557–561 (2019).

20. H. Miwa, R. Dimatteo, J. de Rutte, R. Ghosh, and D. D. Carlo, “Single-cell sorting based on secreted products for functionally defined cell therapies,” Microsyst. Nanoeng. 8, 84 (2022).

21. Y. Yang, S. I. Vagin, and G. Destgeer, “Fabrication of crescent shaped microparticles for particle templated droplet formation,” doi: 10.1101/2023.10.06.561257.

22. G. Destgeer, M. Ouyang, C. Y. Wu, and D. D. Carlo, “Fabrication of 3D concentric amphiphilic microparticles to form uniform nanoliter reaction volumes for amplified affinity assays,” Lab Chip 20, 3503–3514 (2020).

23. G. Destgeer, M. Ouyang, and D. D. Carlo, “Engineering design of concentric amphiphilic microparticles for spontaneous formation of picoliter to nanoliter droplet volumes,” Anal. Chem. 93, 2317–2326 (2021).

24. C. Y. Wu, M. Ouyang, B. Wang, J. D. Rutte, A. Joo, M. Jacobs, K. Ha, A. L. Bertozzi, D. D. Carlo, “Monodisperse drops templated by 3D-structured microparticles,” Sci. Adv. 6 eabb9023 (2020).

25. K. Ha, J. D. Rutte, D. D. Carlo, and A. L. Bertozzi, “Surface energy minimizing configurations for axisymmetric microparticles,” J. Eng. Math. 134, 1–19 (2022).

26. R. S. Du, L. Liu, S. Ng, S. Sambandam, B. H. Adame, H. Perez, K. Ha, C. Falcon, J. D. Rutte, D. D. Carlo, and A. L. Bertozzi, “Statistical energy minimization theory for systems of drop-carrier particles,” Phys. Rev. E 104, 015109 (2021).

27. S. Lee, J. de Rutte, R. Dimatteo, D. Koo, and D. D. Carlo, “Scalable fabrication and use of 3D structured microparticles spatially functionalized with biomolecules,” ACS Nano 16, 38–49 (2022).

